# Costs of dispersal evolution under larval malnutrition

**DOI:** 10.1101/2024.11.17.624046

**Authors:** B. Vibishan, Akshay Malwade, Anish Koner, Vinayak Khodake, Sutirth Dey

## Abstract

Dispersal is a key eco-evolutionary process, and the causes and consequences of its evolution have been well-studied. However, although dispersal depends critically on the nutritional status of organisms, it is unclear how malnutrition affects the evolution of life history and the behavior of organisms under selection for increased dispersal. To address this issue, we used four replicate laboratory populations of *Drosophila melanogaster* previously selected for increased dispersal for 66-67 generations on a protein-poor larval diet. Compared to the unselected controls, the selected populations had lower female body weight, fecundity, male desiccation resistance and mating propensity, but greater male locomotor activity and mating latency. There were no significant differences in aggression. We also compared our results qualitatively with those from a previous study on dispersal selection under a standard larval diet. Barring desiccation resistance, locomotor activity, and rest duration, all other traits investigated responded differently to dispersal selection between the poor and the standard larval nutritional regimes. These results show that the evolution of traits associated with dispersal can differ markedly depending on the nutrition available for the dispersing populations.

## 1 Introduction

In the context of rapid global climate change, predicting the responses of natural populations to changing environments has emerged as an urgent necessity (Kokko and López-Sepulcre, 2006). Dispersal has a well-recognized role in determining the spatiotemporal distribution of populations (Nathan et al., 2008, Kot et al., 1996) and could thus inform how organisms respond to changing environmental conditions. The centrality of dispersal in many ecological and evolutionary processes like population stability, local adaptation, speciation, range expansion etc., has generated much interest in developing ways of predicting the mechanism, pattern, and consequences of dispersal in a wide variety of systems (reviewed in Clobert et al., 2012). However, this is not an easy task as, for an organism, dispersal is a complex process that involves a large number of physiological, morphological, and behavioral traits acting in close coordination. For instance, the process of dispersal typically requires sensory structures that enable decision-making at various dispersal phases (Kautz et al., 2014, Merckx and Van Dyck, 2007, Edelaar et al., 2008) as well as physical features related to facilitating actual movement (Zera and Denno, 1997, Nardi et al., 2008). The energetic demands of inter-patch movement require various physiological processes to ensure sufficient resource availability (Zera and Zhao, 2004).

Similarly, a wide range of behavioral traits, both at the individual and the social level, are liked to dispersal. For example, exploratory movement is an important aspect of leaving the natal patch (Cote et al., 2010, Korsten et al., 2013), while locomotor activity and sampling behavior dictate how organisms collect information about the environment (Kempenaers and Valcu, 2017, Arnold et al., 2017, Hanski et al., 2006). Similarly, aggression is one of a spectrum of social behaviors involved in dispersal initiation and success (Duckworth and Badyaev, 2007, Guerra and Pollack, 2010).

With such a range of traits implicated in the dispersal process, it is likely that a change in, or evolution of, the dispersal phenotype has the potential to change any (or perhaps many) of these traits. At the same time, dispersal evolution can be limited by the inability of an organism to modulate any of these traits for many potential reasons, such as lack of genetic variation, correlated effects of other traits, or environmentally-induced physiological restrictions. For a hypothetical example, suppose an actively dispersing species evolves a greater ability to travel across long distances (dispersal ability *sensu* Tung et al., 2018b) but, at the same time, fails to evolve the tendency to leave a natal patch (dispersal ability *sensu* Tung et al., 2018b). Thus, those organisms that do leave the natal patch can travel long distances, but only a few can initiate dispersal. The distribution of distances dispersed by the members of the population (i.e., the dispersal kernel), and hence the resultant ecological effects of dispersal (e.g., their range expansion rates, settlement success), in such a species would be very different from another species that have evolved both propensity and ability in say, moderate amounts. These ecological effects can potentially modulate how dispersal evolves in these two species.

Traits could thus co-vary with dispersal status in complex manners, and this co-variation could affect how dispersal evolves in a given system. The collection of traits that co-vary with dispersal *en bloc* has been termed the dispersal syndrome (Clobert et al., 2009, 2012) and is an active area of investigation in ecology and evolution across several taxa (Clobert et al., 2012). Previous studies have demonstrated that lab populations of *Drosophila melanogaster* selected for higher dispersal not only evolve higher dispersal capacity (Tung et al., 2018b) but also show correlated changes in a suite of behavioral and metabolomic traits, thus leading to the emergence of a dispersal syndrome (Tung et al., 2018a).

Dispersal syndromes are considered particularly relevant when dispersal traits are difficult to measure directly since measurement of other phenotypic traits as part of the syndrome could be used to predict the dispersal status of the organism. However, this utility of dispersal syndromes for predictive purposes is often equivocal (Sekar, 2012, Lancaster and Downes, 2017). This is because the component traits of a dispersal syndrome can be labile, as different organismal traits can respond to environmental changes to varying degrees (Bowler and Benton, 2005, Vinatier et al., 2011, Jones et al., 2019). The dispersal syndrome in a specific scenario (i.e., a particular population in a given environment) is thus highly context-dependent. Moreover, changes that affect the overall energy budget of an organism could also affect the relative energetic allocation to various traits and thus create the potential for tradeoffs (Losos et al., 2014). Therefore, plasticity in trait correlations and changing energetic budgets with changes in environmental conditions could be important in determining which traits make up the dispersal syndrome and how strong the corresponding associations are.

One of the environmental changes that are likely to make an organism reconsider its energy budget is the availability of food, which is known to be a major driver of dispersal (Fronhofer et al., 2018). However, even when food is available *ad libitum*, changes in the food quality can affect the physiology and life history of the organisms (Prasad and Joshi, 2003, Kolss et al., 2009, Lee and Micchelli, 2013). For example, reduced nutrition can increase the locomotor activity in *Drosophila* (Zúñiga-Hernández et al., 2023), which can potentially increase the short-range dispersal but have a negative effect on long-range dispersal. As mentioned earlier, some previous studies on dispersal evolution (Tung et al., 2018b,*a*, Mishra et al., 2018) have used standard larval nutrition (standard banana jaggery medium; see S2 Text for the recipe). More recently, a similar protocol was used to select for dispersal under a protein-deficient larval nutrition condition (Vibishan et al., 2023). Protein deficiency represents a considerable shift in the larval environment and could potentially change the energetic budget of the organism (Prasad and Joshi, 2003). Therefore, it was unsurprising that these populations that evolved in a protein-deficie(Vibishan et al., 2023). However, this study did not report the associated behavioral and life-history traits accompanying the kernel evolution.

The current study aims to understand whether the trait correlations accompanying dispersal evolution under protein-deficient larval nutrition are qualitatively similar to those that evolve due to dispersal selection under standard nutrition. Accordingly, a range of physiological and behavioral traits are examined in fly populations that have evolved greater dispersal capacity under larval malnutrition and compared to unselected controls that have experienced larval malnutrition but not dispersal selection. The data presented here indicate that selection for dispersal leads to a distinctly different phenotype from the control population in terms of the traits studied here and that the overall dispersal syndrome obtained under larval malnutrition is qualitatively different from that under standard larval nutrition. Several trait correlations with dispersal evolution under larval malnutrition are also markedly different among the sexes, suggesting that dispersal evolution might have very different effects on the males and the females.

## 2 Materials and methods

### 2.1 Ancestral stocks and selection protocol

Detailed descriptions of the maintenance procedure of the ancestral populations have been published elsewhere (Tung et al., 2018a, Vibishan et al., 2023). Briefly, from each of the four ancestral DB*_i_* populations, two populations were derived to give rise to VB*_i_* (Vagabond) and VBC*_i_* (VB-Control). VBs were selected for higher dispersal and VBCs acted as the corresponding controls that did not undergo selection for dispersal. All populations were maintained at a breeding population size of roughly 2400 flies, in discrete generation cycles, under uniform environmental conditions comprising a temperature of 25*^◦^*C and 24h light conditions. In the current study, from each VBC*_i_*, we derived two more populations: MD*_i_* (Malnourished Dispersers) and MC*_i_* (Malnourished Control). Thus, MD*_i_*-MC*_i_* with the same subscript *i* were related by ancestry to the corresponding VBC*_i_*. MDs were selected for higher dispersal using a setup identical to that in the VB selection experiment, while MCs were maintained as corresponding controls not selected for dispersal. The MD-MC populations were raised on a modified banana-jaggery medium with one-third the amount of dry yeast powder as the standard recipe for VB and VBCs. This reduction in yeast corresponds to about 56% lesser dietary protein (see S2 Text for the recipe) and a ∼ 50% reduction in the protein to carbohydrate (P:C) ratio (from 0.17 in the standard diet to 0.08 in the malnourished diet). Eggs from both MD and MC were reared in this protein-deficient diet until eclosion, and the adults underwent selection for dispersal on the 12*^th^* day from egg collection. Initial rearing of stocks was carried out in plastic vials at egg densities of ∼ 30 eggs in about 6mL of food per vial, and subsequently moved to plastic bottles around generation 40, with ∼ 350 eggs in about 60mL of food per bottle.

As with VB-VBC, the selection apparatus consisted of a set of two transparent plastic containers, one of which served as the source and the other as the sink. The source and the sink were connected by a transparent plastic pipe that served as the dispersal path. Neither the source nor the dispersal path contained food or water sources, while a strip of moist cotton was placed in the sink to prevent desiccation after dispersal. On day 12 from egg collection, adult MD flies of both sexes were introduced into the source container and allowed to migrate for a period of 6h, or until about 50% of the flies had reached the sink, whichever occurred first. During this period, the MC flies were kept in a source container (of identical dimension as the MDs), where the opening of the container was closed with a plastic plug. Thus, all the MC flies were allowed to lay eggs for the next generation, which implied that they did not undergo selection for dispersal. The MD and MC adults were transferred to population cages with standard banana-jaggery medium. The adults were provided with live yeast-paste on day 14 and eggs for the next generation were collected on day 15 from the previous egg collection, leading to a 15-day life cycle. The path length used for dispersal selection was increased every few generations to simulate the process of progressively increasing habitat fragmentation. At the start of the experimental evolution, the path length was 2m, whereas by the 83*^rd^* generation (when the last of the assays reported here were performed) the length had increased to 50m. Since MD*_i_*-MC*_i_* with the same subscript were related by ancestry, they were assayed together and considered a single block in all statistical analyses. All assays were performed after one generation of common garden rearing in the larval malnutrition diet but without dispersal selection to minimize the effects of non-genetic parental effects due to dispersal selection. In order to avoid larval crowding, eggs for common garden rearing were collected at a density of about 350 per bottle containing about 60mL of food, and for the assays, about 30 eggs per vial containing about 6mL of food.

Assays to measure various physiological and behavioral traits were performed to elucidate if selection for dispersal under larval malnutrition had led to correlated responses between different aspects of life history. In all cases, assays were performed two blocks at a time due to logistic constraints.

### 2.2 Physiological assays

Dry body weight was measured as a proxy for body size in generation 66-77. 12 days post egg collection, flies were sorted by sex under mild CO_2_ anesthesia, and then placed in 1.5mL Eppendorf tubes, with 20 individuals per tube and 10 such replicates for each sex and population. The flies were killed by flash freezing in liquid nitrogen and dried in the tubes at 60*^◦^*C for 72h. Following this, the weight of each Eppendorf tube was measured to the closest 0.1mg on a digital weighing balance, first with and then without the flies. The average body weight per fly was calculated as: 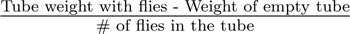.

Fecundity was measured as the number of eggs laid per male-female pair in each population in generation 82-83, assayed two blocks at a time. On day 14 post egg collection, 50 male-female pairs were collected under mild CO_2_ anesthesia and each pair was placed in a 50mL Falcon tube with holes for ventilation. The lid contained a small food cup that served as a flat surface for oviposition. These set-ups were left undisturbed for 14 hours in a well-lit environment maintained at 25*^◦^*C. After 14h, the flies were discarded and the number of eggs laid on the food cup was counted under a stereomicroscope.

Development time was measured in generation 66-67 as the time taken for eclosion from the time of egg collection. Each population was given an egg-laying period of about 14h, after which exactly 30 eggs were collected and placed in vials containing about 6mL of the malnutrition food. 10 replicate vials were set up for each population and the vials were monitored for eclosion starting from around day 8 after egg collection, once pupae had started to darken. Freshly eclosed flies were subsequently collected every 2h under mild CO_2_ anesthesia, and the numbers of males and females emerging at each time point were recorded.

Desiccation resistance was measured in generation 66-67 as the time it took for a fly to die due to lack of food and water. For each population, same-sex groups of 10 flies each were collected on day 12 from egg collection under mild CO_2_ anesthesia and placed in fresh glass vials without any food or moisture. 10 such replicate vials each of males and females were set up for each population, and monitored every 2h. At each time point, the number of flies alive was recorded.

### 2.3 Behavioral assays

Locomotor activity and resting behavior of adult male flies was assayed in generation 66-67 using the Drosophila Activity Monitoring (DAM2) data collection system (Trikinetics Inc., Waltham, MA) following standard protocols (Chiu et al., 2010, and Text S1.2.1 therein). On day 12 from egg collection, individual males were orally aspirated singly into small glass tubes (length 6.5cm, diameter 5mm), with 32 replicates per population. These tubes were placed in the monitoring apparatus such that two perpendicular IR beams pass through the center of each glass tube. Activity was recorded for 6h after a 15 minutes period for acclimation. The activity for a given fly was estimated as the average number of times the fly crossed the IR beam per hour (Chadov et al., 2015). Continuous inactivity for five minutes or more was considered as sleep/rest and the amount of time spent in rest was calculated as a proportion of the total recording duration (Hendricks et al., 2000, Chiu et al., 2010).

Aggressive behavior was studied in generation 66-67 by means of male-male fighting bouts between MD and MC within each block (e.g. MD_1_ male against MC_1_ male and so on). Following published protocols (Yurkovic et al., 2006), approximately 30 eggs were collected from each population and placed in vials containing about 6mL of the malnutrition food. The first few flies to eclose were discarded to collect flies from the eclosion peak and the vials were subsequently checked once every 6h (Yurkovic et al., 2006). As fruit flies take about 8h from eclosion to become sexually active, a 6h monitoring period would ensure that the flies collected would be virgins (Prasad and Joshi, 2003). Flies eclosing in each 6h interval were collected under mild CO_2_ anesthesia and the males were isolated singly in food vials with about 2mL of food until the day of the assay. This isolation was necessary to prevent social contact with other males as this could lead to the formation of hierarchies through aggressive interactions and affect their performance during the assay (Chen et al., 2002, Kravitz and Fernandez, 2015). About 24h before the assay, the males from both populations were exposed to daylight fluorescent pigments (DayGlo) by loading a small amount of the pigment powder on the cotton plug of the vial. This coated the fly in the corresponding color, enabling easy identification. On day 12 from egg collection, arenas were assembled in 12-well clear plastic plates (Corning) with a piece of standard banana jaggery food in the center of the well. A decapitated female from an unrelated baseline population was stuck to the center of the food piece with yeast paste and one male of each population was introduced into the arena by aspiration. The food piece and the decapitated female acted as proximate cues to induce the males to fight (Yurkovic et al., 2006). The arenas were recorded using video cameras for a period of 1 hour under well-lit conditions. Since we needed to use two color pigments for distinct identification of MD vs. MC flies within each arena, both populations were assayed in both color schemes to control for any possible biases in behavior due to the specific coloring. For each block therefore, 18 replicate arenas were assayed in which the MD male was loaded with a green pigment and the MC male with pink, and 18 more replicate arenas were assayed in which the MD male was pink and the MC male was green. This lead to 36 replicates of MD-MC fights in each block, 18 in each color scheme. Aggression behavior was scored in terms of victory in fighting bouts, after disregarding the first 5 minutes due to acclimation. Following previous studies (Yurkovic et al., 2006), a given fly was considered the winner of a fight if it successfully chased the other fly out of the food piece three times in a row.

Mating traits were measured in generation 66-67. Following previous studies (Syed et al., 2020, Maggu et al., 2021), virgin males and females were collected for each population from the eclosion peak as described above and isolated in same-sex vials of 8 flies each with about 2mL food. On the day of the assay, one male and one female were introduced into a vial containing about 2mL of 1.3% agar and their interactions were recorded on video for a period of 1h. For each population, 24 such replicate vials were recorded. The solidified agar allowed for the mitigation of desiccation stress. From the recordings, the times when mating began and ended were noted for each vial in which mating occurred within the observed period. Three metrics were calculated from this data to characterize the mating properties of each population. Mating latency, i.e., the time taken for initiation of mating, was calculated as the difference between the start times of mating and the video time when the vial was first introduced into the recording arena. Copulation duration was calculated as the difference between the mating start and end times, and mating propensity was calculated as the proportion of male-female pairs that mated during the period of observation.

### 2.4 Statistical analyses

Except for the aggression data, all other statistical analyses were carried out using general or generalized linear mixed-effect models with the lmerTest package (Kuznetsova et al., 2017) in R (R Core Team, 2021, version 4.1.2). Dry body weight was analyzed in a mixed-effects general linear model (GLM) with Gaussian error distributions, considering selection and sex as fixed effects, and block as a random intercept. Average activity counts per hour, mating latency and copulation duration were likewise analyzed in separate Gaussian mixed-effects GLMs with selection as a fixed factor and block as a random intercept. Rest fraction and mating propensity were both calculated as proportions, and were therefore analyzed using mixed-effects generalized linear models (GLMs) with binomial errors and a logit link function, with selection as a fixed effect and block as a random intercept. For rest fraction, the response variable consisted of the total duration spent in rest defined as success and the total active duration defined as failure. For mating propensity, the response variable consisted of the number of pairs that mated defined as success and the number that failed to mate defined as failure. For the time to desiccation (10 adults per vial) and time to eclosion (30 eggs per vial) assays, we analyzed the vial means using mixed-effects Gaussian GLMs. Time to eclosion was calculated from the time of assay setup on the day of egg collection up to whenever each fly eclosed. For these GLMs, selection and sex were considered fixed effects, with block as a random intercept. Significance of individual effects or interactions from each model was inferred based on *χ*^2^ statistics from analysis of deviance at the 5% significance level. In all cases, further comparisons of group means within the model were carried out using pairwise differences from the emmeans package (Lenth, 2023, version 1.8.4-1) and Cohen’s *d* was calculated using appropriate functions from the psych package to estimate effect sizes (Revelle, 2022, version 2.2.9). All group mean comparisons and effect sizes were calculated as the MD-MC difference, and could be positive (MD *>* MC), negative (MD *<* MC) or zero. The absolute values of effect size were therefore interpreted as large, medium and small for |*d*| ≥ 0.8, 0.8 *>* |*d*| *>* 0.5, and |*d*| *<* 0.5 respectively (Cohen, 2013).

The response variable from the aggression assay was the number of wins for a given population. This was pooled across the two color schemes, and compared between MD and MC using a two-sided Wilcoxon’s rank-sum test using the wilcox.test() function from the stats package (R Core Team, 2021, version 4.1.2).

## 3 Results

This study investigates the consequences of dispersal selection in the background of a protein-deficient larval diet. Earlier studies have elucidated the effects of dispersal selection under standard nutrition (Tung et al., 2018b,*a*), as well as a marked difference in the evolution of dispersal capacity under larval malnutrition (Vibishan et al., 2023). Malnutrition could alter the allocation of resources across various life processes and thus potentially change the overall patterns of trait correlations and trade-offs. This is examined here by measuring a suite of physiological and behavioral traits in fruit fly populations selected for increased ambulatory dispersal, and comparing their trait values with those of unselected control populations. All relevant statistical information has been reported inline when relevant, and S1 Text has the details of all the GLMs and subsequent analyses.

### 3.1 Physiological trait responses to dispersal selection are sexually dimorphic

Figure 1 shows a summary of all responses in physiological traits because of dispersal selection under larval malnutrition. While sexual dimorphism in adult fly body weight is well-known (Prasad and Joshi, 2003), we also found a significant selection × interaction in the mixed-effects GLM (*χ*^2^ = 30.11, *p <* 0.0001; Figure 1A). The average body weight of the females of the dispersal-selected populations (MDs) was significantly less than those of the controls i.e., MCs (Cohen’s *d* = −1, t_153_ for mean difference = −8.838, *p <* 0.0001). However, there was no significant difference among the males (*d* = −0.19, t_153_ for mean difference = −1.078, *p* = 0.2827). Adult body weight in flies could affect fecundity through the total available biomass (Hoňek, 1993, Prasad and Joshi, 2003, Tu and Tatar, 2003). Figure 1B suggests that fecundity is lower in MD than MC, and the corresponding mixed-effects model showed a significant effect of selection (*χ*^2^ = 4.182, *p* = 0.0408). While this observation aligns with the difference in female body weight between MD and MC, the effect size corresponding to the difference in fecundity is rather small (*d* = −0.06). This could therefore be an emerging trend that may progressively diverge with further selection.

**Figure 1:**
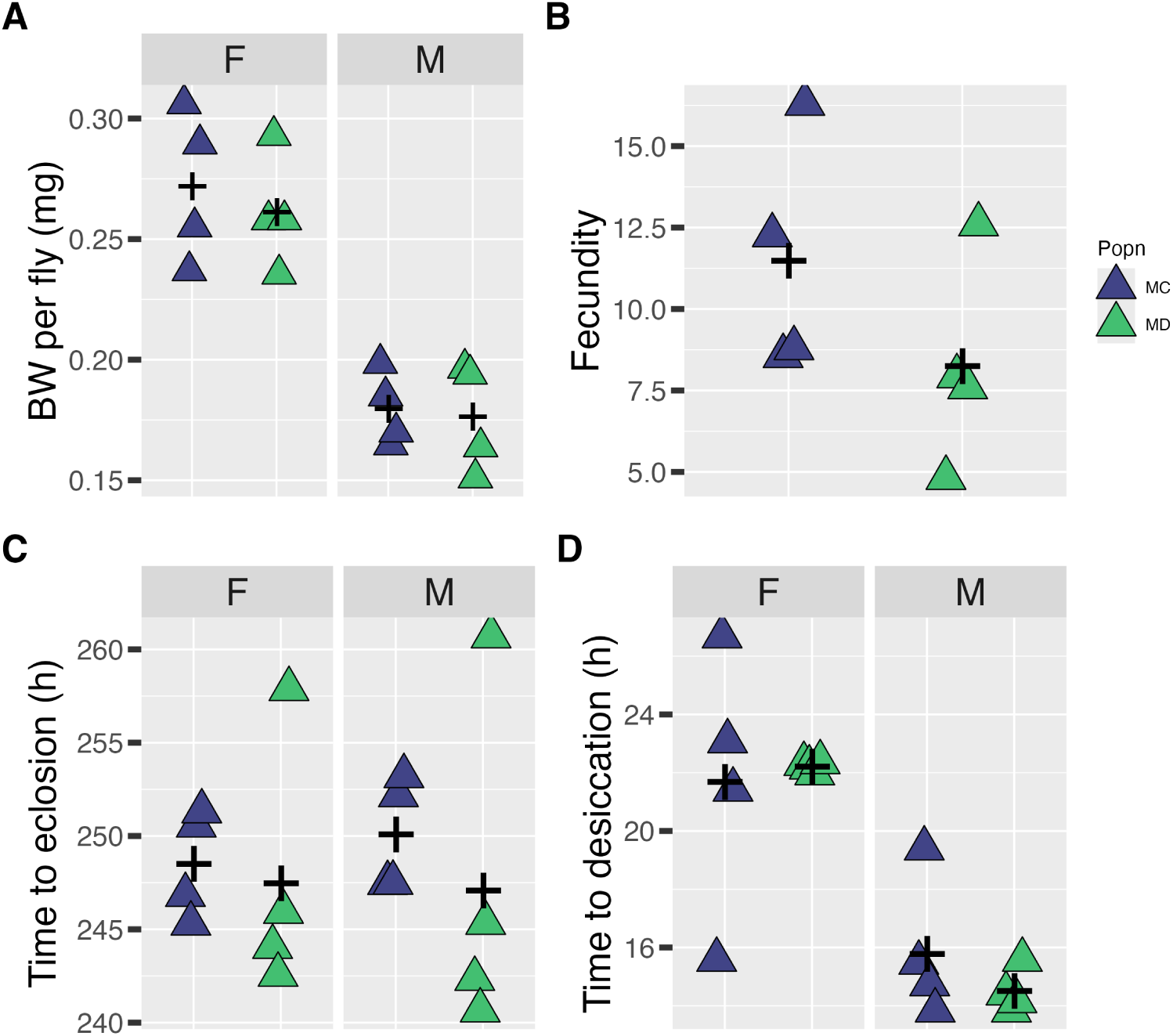
Physiological trait responses due to dispersal selection under larval malnutrition. (A) dry body weight measured in generation 66-67, except for block 2 which was measured in generation 77; (B) fecundity measured as clutch size per mating pair in generation 82-83; (C) desiccation resistance measured in generation 66-67 as the time taken to death under desiccation; and (D) developmental time measured as the time period between egg collection to adult eclosion in generation 66-67. All points are average values within each block and the black plus sign is the average of the block means.

Time to death by desiccation shows a distinctly different MD vs MC trend among the sexes (Figure 1C). Both the main effect of sex (*χ*^2^ = 109.99, *p <* 0.0001) and the selection × sex interaction are significant (*χ*^2^ = 5.41, *p* = 0.02). The main effect of sex is not surprising as *Drosophila* females are heavier and could have greater biomass to survive longer under desiccation (Clark and Doane, 1983, Graves et al., 1992). However, the MD females show a trend towards longer time to desiccation, although the difference is not statistically significant, with only a modest effect size (*d* = 0.16, t_152_ for mean difference = 0.926, *p* = 0.3558). Among males, MDs died significantly faster under desiccation than MC (*d* = −0.61, t_152_ for mean difference = −2.359, *p* = 0.0196).

Finally, patterns in development time were complicated (Figure 1D). The mixed-effects GLM reveals no significant effect of selection or sex, and the selection × sex interaction is not statistically significant (*χ*^2^ = 1.529, *p* = 0.2163). However, pairwise comparison of group means within each sex reveals a significant difference in time to eclosion between MD and MC only in males, albeit with a small effect size (*d* = −0.41, t_153_ for mean difference = −2.556, *p* = 0.0116). Females show an even smaller effect size and insignificant mean difference in their development times (*d* = −0.15, t_153_ for mean difference = −0.807, *p* = 0.4208). Visual inspection of Figure 1D suggests that there could be a considerable difference in the mean time to eclosion between MD and MC, particularly in the males, but one point each in MD males and females appears to be an outlier, which corresponds to the mean time for block 4. Removing this point from the analysis leads to a significant selection × sex interaction (*χ*^2^ = 7.549, *p* = 0.006) and large effect sizes for the MC vs MD difference in time to eclosion (*d* = −1.3, t_134_ for mean difference = −4.422, *p <* 0.0001 for females; *d* = −1.9, t_134_ for mean difference = −8.8494, *p <* 0.0001 for males). Therefore, male MDs could be developing faster than MC males, while a similar trend may be emerging in the females.

### 3.2 Behavioral correlates of dispersal evolution under larval malnutrition

Average activity levels are clearly elevated in MD relative to MC (Figure 2A) with a significant selection effect (*χ*^2^ = 109.88, *p <* 0.0001) and a large effect size (*d* = 1.27, t_251_ for mean MC vs MD difference = 10.482, *p <* 0.0001). Likewise, Figure 2B shows that the rest fraction, which the proportion of the monitoring period spent in a rest state, is considerably reduced in MD relative to MC (*χ*^2^ = 6352.72, *p <* 0.0001, MD/MC odds ratio = 0.1312, *z* = 79.704, *d* = −0.84). These two observations reveal a more active, restless locomotor phenotype in MD relative to MC due to dispersal selection, similar to what was seen earlier due to dispersal selection under standard nutrition (Tung et al., 2018a).

**Figure 2:**
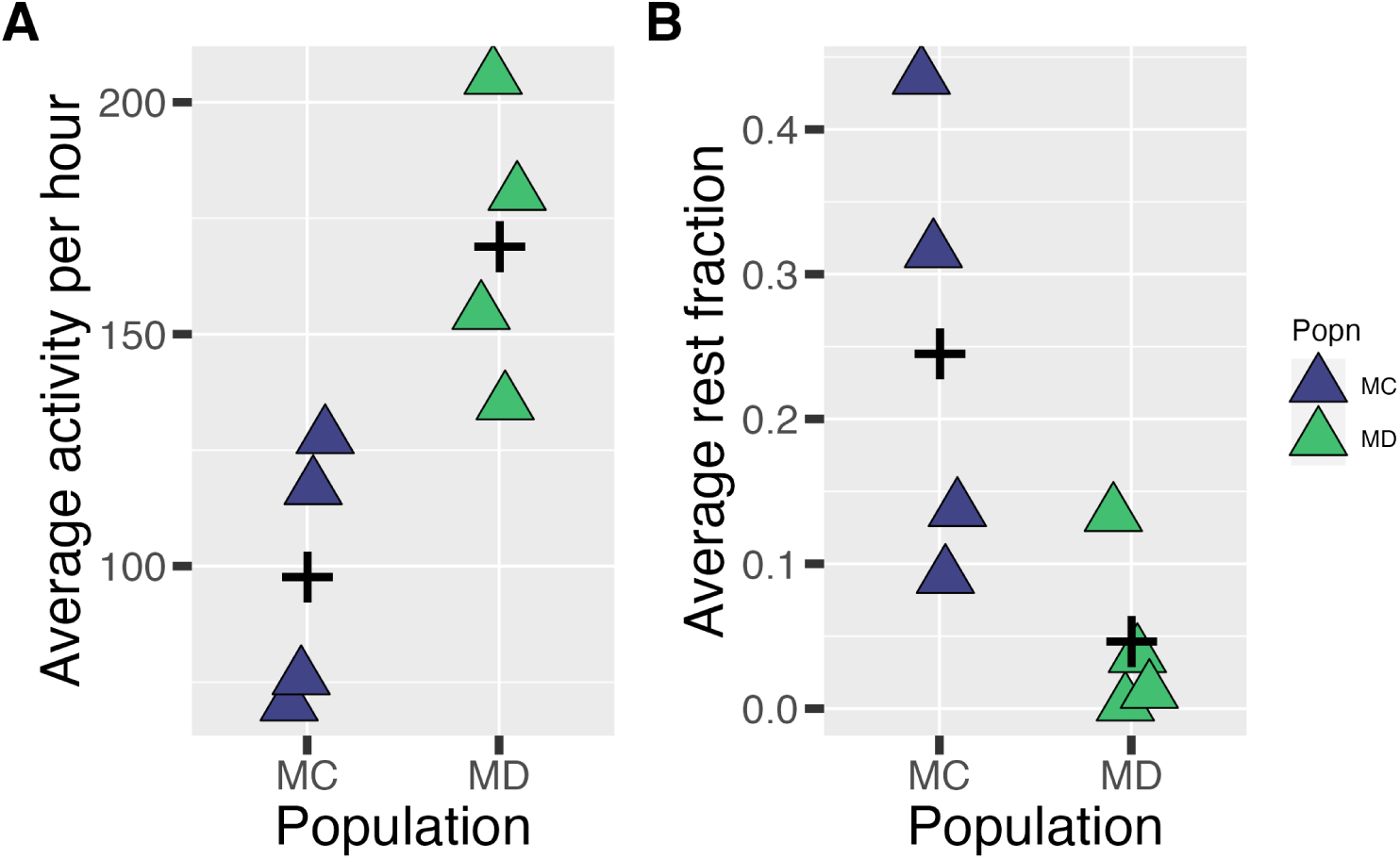
Locomotor phenotype under dispersal selection. Changes in (A) activity counts per hour and (B) fraction of rest duration due to dispersal selection under larval malnutrition; all measurements were made after 66-67 generations of selection. All points are average values within each block and the black plus sign is the average of the block means.

The average mating latency of the MDs was not different from that of MCs (Figure 3A; *χ*^2^ = 0.0901, *p* = 0.764); pairwise comparisons of the group means produced a very small effect size (*d* = −0.05, t_91.6_ for mean difference = 0.286, *p* = 0.016). On the other hand, MDs had a significantly shorter copulation duration (*χ*^2^ = 5.794, *p* = 0.016) even though the corresponding effect size is small (*d* = −0.39, t_155_ for mean difference = −2.404, *p* = 0.0174; Figure 3B). Mating propensity, defined as the fraction of pairs that mated during the experiment, shows a significant effect of selection (*χ*^2^ = 4.279, *p* = 0.0385) and a comparison of the group means shows that the odds of mating propensity for MD (MD/MC odds ratio = 0.435, *z* = −2.069, *p* = 0.0389, *d* = −1.15). Finally, the average number of fights won by MC vs. MD is similar, and a Wilcoxon’s rank-sum test confirms that the two populations do not differ in their winning tendencies (W = 12.5, *p* = 0.2425; Figure 4).

**Figure 3:**
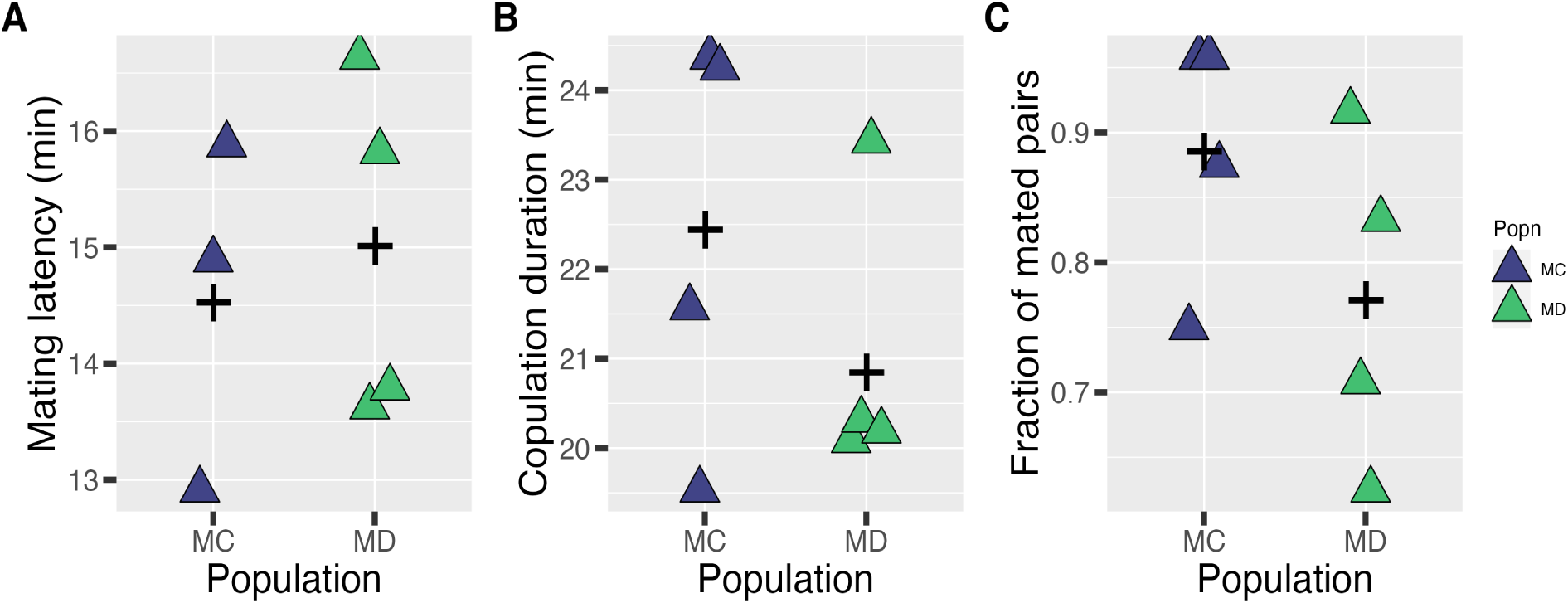
Impairment in mating traits due to dispersal selection. Changes in (A) mating latency, (B) copulation duration and (C) mating propensity between MD and MC; all measurements were made after 66-67 generations of selection, and all points are average values within each block and the black plus sign is the average of the block means.

**Figure 4:**
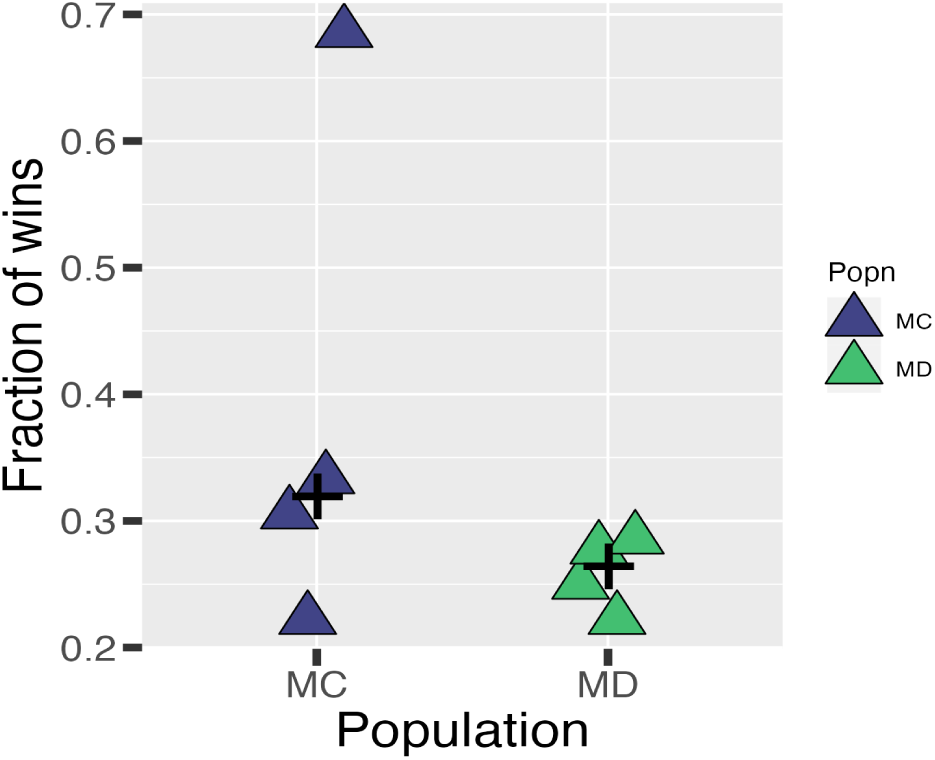
Aggressive behavior is not associated with dispersal evolution. The figure shows the fraction of chase-aways between MD and MC. All measurements were made after 66-67 generations of selection, and all points are number of wins within each block. The black plus sign is the median number of wins for each population across the four blocks.

## 4 Discussion

### 4.1 Dispersal selection under malnourished conditions affects body weight and fecundity

The dry body weight of the dispersal-selected (MD) females to be lower than those of the control (MC) females (Figure 1A). As dry body weight is typically used as a proxy for body size and, to some extent, body condition (Knapp, 2016), the reduction in body weight could imply that the MD females are showing signs of a cost of dispersal selection. An association between body size and dispersal has been documented across taxa, with or without distinct dispersal polymorphisms (Zera and Denno, 1997, Braendle et al., 2006, Jenkins et al., 2007, Asplen, 2018). In general, body size is expected to be positively associated with dispersal, either through mechanical improvements in the efficiency of locomotion (Nardi et al., 2008, Lancaster and Downes, 2017, Berwaerts et al., 2006, Ohlberger et al., 2006) or the availability of greater energetic reserves (Reim et al., 2019). On the other hand, a negative association of body size with active dispersal has been found in some cases in which a larger or heavier body increases the energetic cost of locomotion (Dahirel et al., 2021). The net association between body size and dispersal in active dispersers can thus be seen as the resultant of the extent to which a larger body size leads to potentially greater energetic reserves instead of increased energetic demands. Lower body weight among female dispersers in our study could be indicative of the latter case, where large body size under a poor nutrition regime might be a substantial energetic demand for dispersers. In turn, this could represent a direct cost of dispersal evolution.

The reduction in body weight in the MDs is consistent with their lower fecundity (Figure 1B), as the number of eggs laid is generally expected to be positively correlated with female body weight in *Drosophila* (Prasad and Joshi, 2003). Similar to body weight, lower fecundity in MD could indicate a further cost of dispersal evolution. Resource allocation between dispersal and fecundity can differ between range margins and cores, favoring dispersive strategies at margins at the expense of reproductive investment (Hughes et al., 2003). More broadly, lower fecundity in the dispersers in our study is consistent with a potential flight-oogenesis syndrome, which signifies the reduction and/or postponement of reproductive investment *vis-à-vis* dispersal (Roff, 1990). Although closely associated with wing dimorphism in migrating insects (Mole and Zera, 1993, Zera and Denno, 1997, Lorenz, 2007), temporal separation between reproduction and dispersal has also been recorded in monomorphic migrant butterflies (Stefanescu et al., 2021). Hormone-signaling based modification of metabolism is a common mechanistic link between dispersal and reproduction for wing-dimorphic insects like crickets (Zera and Zhao, 2004) and paper wasps (Southon et al., 2020). However, corresponding metabolic or physiological processes underlying a dispersal-reproduction tradeoff in monomorphic insects are yet to be identified. Given that female dispersers in this study show a trend consistent with such a trade-off (Figures 1A and B), this offers a suitable experimental system in which to address the corresponding physiological mechanisms.

## 5 Emerging sexual dimorphism due to dispersal selection

The quality of larval diet is known to affect the sexes differently in *Drosophila melanogaster*. For example, in a protein-rich diet, the body size of females, but not males, is positively affected (Millington et al., 2021). This is consistent with the general observation from one-generation dietary manipulation studies in *Drosophila* that traits in females are typically more sensitive to nutritional changes than in males (Shingleton et al., 2017, Lee and Jang, 2014). In our experiments, the selection for dispersal was playing out in the background of reduced protein availability, which could have affected the sexes differently. Moreover, due to our selection protocol, there is a difference between the nature of the selection on the males and the females. Since oviposition for the next generation happens exclusively after dispersal, female reproductive success in our experiments is contingent on reaching the destination patch. On the other hand, male reproductive success could be realized through mating at any point after reaching sexual maturity, regardless of whether those mating opportunities arise before, during or after the dispersal event. Thus, selection pressure on males to reach the destination patch successfully is likely weaker than that on females. However, there is an added layer of complexity here. The strategy for abandoning dispersal in favor of mating will also require that the corresponding males evolve strong sperm defenses to ensure that males who mate later (i.e., after dispersal) do not get a significant advantage in reproductive fitness (last male advantage; Singh et al., 2002, Schnakenberg et al., 2012). This is why we expect at least some sexual dimorphism in terms of the evolved traits, although it is difficult to predict the precise traits in which this dimorphism might manifest.

Although MD males do not differ from MC males in terms of their dry body weight (Figure 1A), they do show a trend towards lower desiccation resistance (Figure 1D). This suggests that in males, unlike females, the cost of dispersal evolution could manifest in terms of desiccation resistance rather than body weight. While body weight is generally positively associated with stress tolerance (Clark and Doane, 1983, Arnett and Gotelli, 2003, Knapp, 2016), desiccation resistance, in particular, is closely related to the total body levels of glycogen rather than the actual biomass (Graves et al., 1992). Thus, MD and MC males may differ in terms of their glycogen content without a significant difference in net biomass, which (if true) would explain the difference in desiccation resistance between males of the two populations. A possible explanation for the evolution of a male-specific desiccation resistance could be differences between the strength of selection pressure for dispersal among the two sexes. As discussed in the last paragraph, the males face a somewhat weaker selection pressure to reach the destination patch than the females, which might reduce the need for improved desiccation resistance in the former. Interestingly, previous data have shown a lower tendency to initiate dispersal in MD males than MD females (Vibishan et al., 2023). Taken together, these trends indicate that the sexes have responded differently to dispersal selection.

Finally, our data indicate that there were no effects of selection for dispersal on the development times of either sex (Figure 1C). However, one population (MD4) drove this lack of effect. When we remove the data for that one population, dispersers of either sex tend to develop faster than non-dispersers (see S1 Text for details). Within the context of a so-called pace-of-life syndrome, higher dispersal is expected to be associated with a faster growth rate (Ŕeale et al., 2010), which is consistent with the developmental time responses in both sexes in our study. In general, adult body weight is known to be positively associated with the time taken for eclosion in Drosophila (Prasad and Joshi, 2003, Tu and Tatar, 2003). Therefore, all else being equal, our MD flies are expected to be lighter than the MC flies. This prediction was borne out in the females, but not males (Figure 1A). On the other hand, while females in our study seem to show a cost of dispersal evolution through body weight, males seem to manifest the cost of dispersal in terms of desiccation resistance. This indicates that dispersal selection has led sex-specific physiological responses. In this context, it is important to note that whether or not a faster growth rate leads to lower adult body weight is known to be dependent on the development phase in which growth acceleration has taken place (Vijendravarma et al., 2012). Therefore, disperser males and females could have evolved growth acceleration in different developmental phases. This, in turn, could involve different sets of physiological mechanisms operating in a sex-specific manner, leading to sex-specific costs of dispersal. If substantiated, this would add to a growing pool of knowledge regarding sexual dimorphism in dispersal evolution (Mishra et al., 2018, Li and Kokko, 2019).

### 5.1 Behavioral correlates of dispersal indicate a potential mechanism?

Since our dispersal selection involved ambulatory movement, locomotor activity is one of the most direct indicators of the response to selection. As expected, the dispersal selected populations had greater locomotor activity over a 6h period than unselected controls (Figure 2A). Likewise, the fact that selected flies tend to rest for a smaller part of the total duration (Figure 2B) points to a more active and restless behavioral phenotype.

In fruit flies, dopamine activity is a key mediator of increased locomotor activity and reduced sleep (Kume et al., 2005, Sitaraman et al., 2015). Neuronal processes surrounding dopmaine signaling in the fruit fly are complex, involving multiple classes of excitatory and inhibitory receptors implicated in various functions (Zhang et al., 2016). Recent evidence has shown that poor larval nutrition can significantly increase adult activity by upregulating dopaminergic neuronal pathways (Zúñiga-Hernández et al., 2023). This is consistent with personal observations, which indicate that average levels of locomotor activity are generally elevated in both MCs and MDs relative to other fly populations raised on standard nutrition (unpublished data). Therefore, it is tempting to speculate that the increased activity in the MD flies might be driven by increased levels of dopamine. However, a broader scrutiny of the trends reveals that the situation might be more complicated.

In the MD flies, higher locomotor activity is seen to be associated with no changes in mating latency (Figure 3A), but a significant reduction in copulation duration (Figure 3B) and mating propensity (Figure 3C). On the other hand, increased dopamine levels in adult fruit flies are expected to enhance mating receptivity in females (Neckameyer, 1998) and courtship behavior in males (Lim et al., 2018, Love et al., 2023), both of which (singly or in combination) should lead to a reduction in the mating latency and an increase in the mating propensity. Thus, the trends in the mating-related traits do not match with those expected from an increase in dopamine levels. However, this discrepancy is ameliorated when we take into account that in *Drosophila*, inducing higher dopamine signaling within the first few hours of adult eclosion leads to impaired mating behavior later in life, around days five to twelve from eclosion (Kayser et al., 2014). Selection for dispersal in our system happens within 2-3 days from eclosion, and the locomotor activity of the flies is high at this time (Figure 2A). This suggests that dopamine activity could also be elevated during the same time, which (if true) could explain impaired mating in the MDs. However, it remains unclear if the phase of higher locomotor activity identified in our flies corresponds sufficiently to the critical time period of early adult life in which the importance of dopamine signaling has been demonstrated (Kayser et al., 2014).

In summary, our current results suggest that dopamine might be an interesting starting point for elucidating the mechanistic basis of dispersal evolution under larval malnutrition. However, since we have yet to measure dopamine levels directly, we cannot make definitive statements on this aspect.

### 5.2 Dispersal “syndromes” under standard vs. poor larval nutrition

Finally, we examine the findings of this study in the context of an earlier work on the evolution of dispersal under standard nutrition in *Drosophila melanogaster* (Tung et al., 2018a, Mishra et al., 2018, 2022). The two studies used the same dispersal set up, environmental conditions like temperature and humidity, and mostly assayed similar traits. Although this implies that any differences between the qualitative results of these two studies are likely to be attributable primarily to the differences in the nutritional regimes, such across-study comparisons have some potential limitations. Responses to selection are inherently variable, and given other logistic constraints, phenotypic data under the two nutrition regimes were collected over a range of generations. Since corresponding phenotypes cannot be measured in both nutritional regimes in precisely the same generation, quantitative comparisons between the two datasets are not valid. Also, since the two studies are separated by approximately six years, the life-history data were collected at different times. This is why we avoid putting the two datasets into a common statistical analysis. Instead, we limit ourselves to comparing qualitative trends emerging over a more extended period, which might indicate overall trends in dispersal evolution. Given the difference in the nutritional background, such a comparison could shed light on the context dependence of dispersal evolution and its correlates.

Table 1 summarizes behavioral and physiological traits whose correlated evolution with dispersal were compared between standard (Tung et al., 2018a) and poor larval nutrition (this study). It is clear that with the exception of desiccation resistance, locomotor activity and rest duration, all other traits have responded differently to dispersal selection between the two nutritional regimes (note that development time is not included in this table as comparable measurements were not available from the standard nutrition populations). Taken together, there are clear indications that protein deficiency through the larval stages represents a sufficiently severe stress for several trade-offs to develop regarding physiological traits. The shift in the difference in male-male aggression between MD and MC, apart from being an additional tradeoff of dispersal, could be of interest regarding the mechanism of dispersal evolution under larval malnutrition. The flies selected for dispersal under standard nutrition had elevated levels of octopamine, which was thought to be the reason for their increased aggression (Tung et al., 2018a). The lack of a similar increase in aggression in the MD flies (Figure 4) suggests that octopamine signaling may not be as involved in dispersal evolution under malnutrition.

**Table 1:** Comparing trait correlations of dispersal evolution under varying conditions of larval nutrition. . The second and third columns describe the change in each trait in the disperser against the non-dispersers; lower therefore implies that the trait value was lower in the dispersers than the non-dispersers, and so on. The numbers in parentheses indicate the generation numbers over which data were collected in each case. Poor nutrition studies were carried out in the MD-MC populations (second column), while standard nutrition studies were carried out in the VB-VBC populations (third column), as published elsewhere (Sruti, 2017, Mishra et al., 2018, Tung et al., 2018a). *^†^*Only measured in males.

| Trait | Poor nutrition<br>(66-83) | Standard nutrition<br>(49-70) |
| --- | --- | --- |
| Dry body weight | Lower in females | No difference |
| Early-life fecundity | Lower | No difference |
| Desiccation resistance | Lower in males | Lower |
| Locomotor activity | Higher<sup>†</sup> | Higher<sup>†</sup> |
| Rest duration | Lower<sup>†</sup> | Lower<sup>†</sup> |
| Male-male aggression | No difference | Higher |
| Mating characteristics | Mating latency higher |  |
|  | Copulation duration lower | No difference |
|  | Mating propensity lower |  |

## 6 Conclusions

Predicting dispersal outcomes is typically difficult due to the high context specificity of the process (Vinatier et al., 2011, Travis et al., 2013, Legrand et al., 2016, Jones et al., 2019). Studying its evolution is even more challenging as we know that changing the environmental context can alter the outcomes of evolution, even if the nature of the selection remains the same (Burgess et al., 2016, Reim et al., 2019). To complicate matters further, it can be challenging to predict evolutionary changes based on results from one-generation experiments, particularly in the context of dispersal evolution (Mishra et al., 2018). That is why, although an earlier study had found dispersal could evolve without any major life-history costs under standard nutrition (Tung et al., 2018a), there was no easy way to predict the outcomes under slightly less ideal scenarios like slight malnutrition. In the current study, the amount of larval malnutrition was enough for the manifestation of costs of dispersal, where none (except a slight decrease in desiccation resistance) were found under standard nutrition. This is reflected in the reduction in female body weight, fecundity, and mating propensity in the selected populations of the current study. Given that, in nature, many populations probably face protein shortages in their diet, our current study suggests that such populations are likely to pay costs for the evolution of greater dispersal tendencies. This will limit the extent to which dispersal (and its associated syndrome) will evolve in such populations, which in turn might alter how they affect or get affected by other ecological processes like range expansion or colonization of a new habitat (Bowler and Benton, 2005, Nathan et al., 2008). Thus, the search for a predictive framework for the effects of dispersal continues.

## Acknowledgements

The authors are grateful to Aarcha Thadi, Subhashree Subhadarshini, Rakesh M, Pranjali Singh, Pratik Joshi, and Ruchitha B G for their support with stock maintenance and data collection. The authors acknowledge Nikita Sabnis for initiating the MD-MC selection experiment on which the study is based.

## S1 Text for “Costs of dispersal evolution under larval malnutrition”

### Statistical analyses

For each assay, details of statistical analyses are given in three parts:

1. The code used to fit the GLM is reproduced, followed by the analysis of deviance table from the corresponding model object. All inferences of statistical significance are based on the test statistics in this table.
2. Pairwise comparisons for each trait from the corresponding model object.
3. Cohen’s *d* for each dataset

In this study, data from the male-male aggression experiment were compared using a two-sample Wilcoxon’s signed rank test, and therefore does not involve analyses of the type given above.

## S1 Physiological traits

### S1.1 Dry body weight

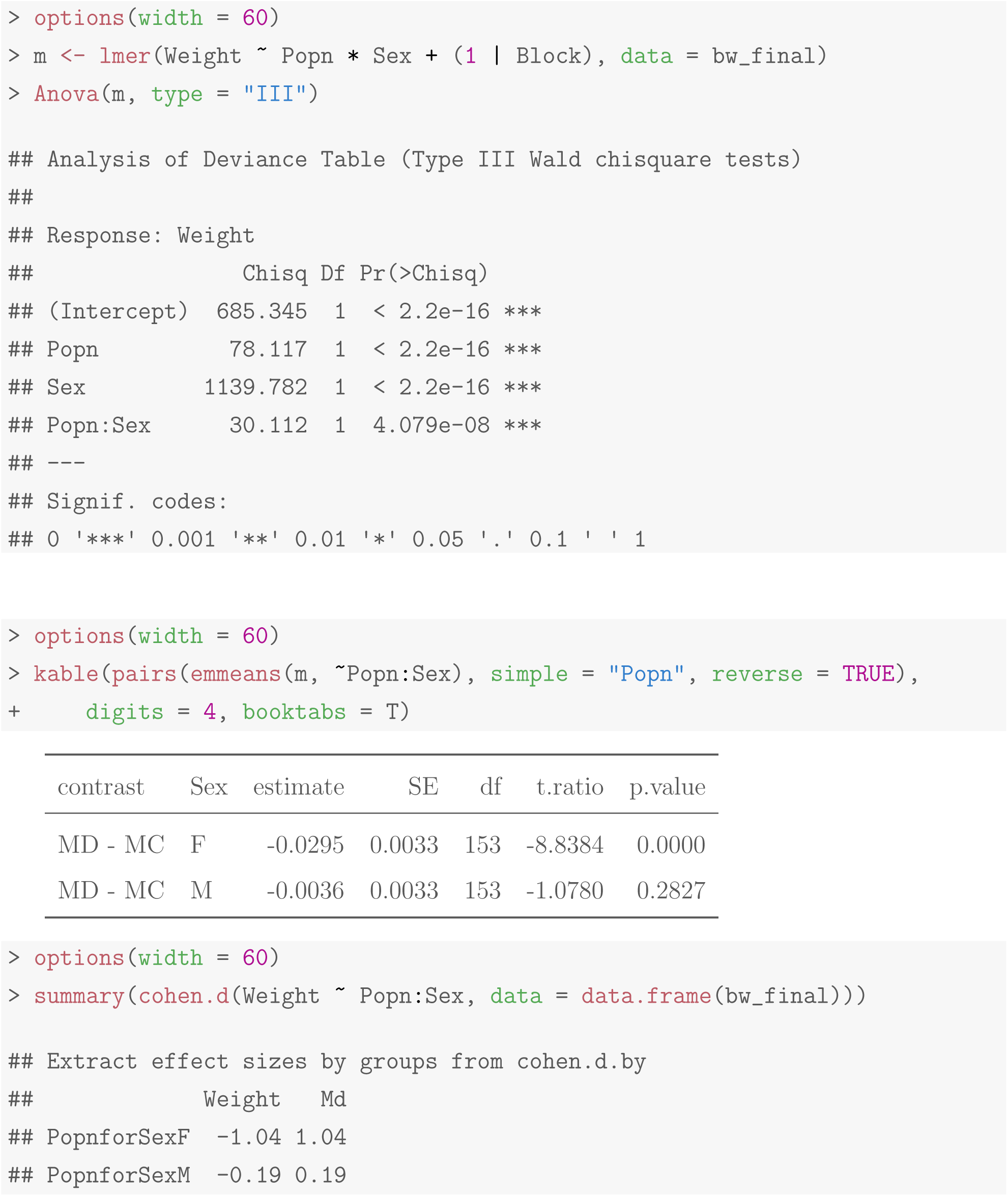

### S1.2 Fecundity

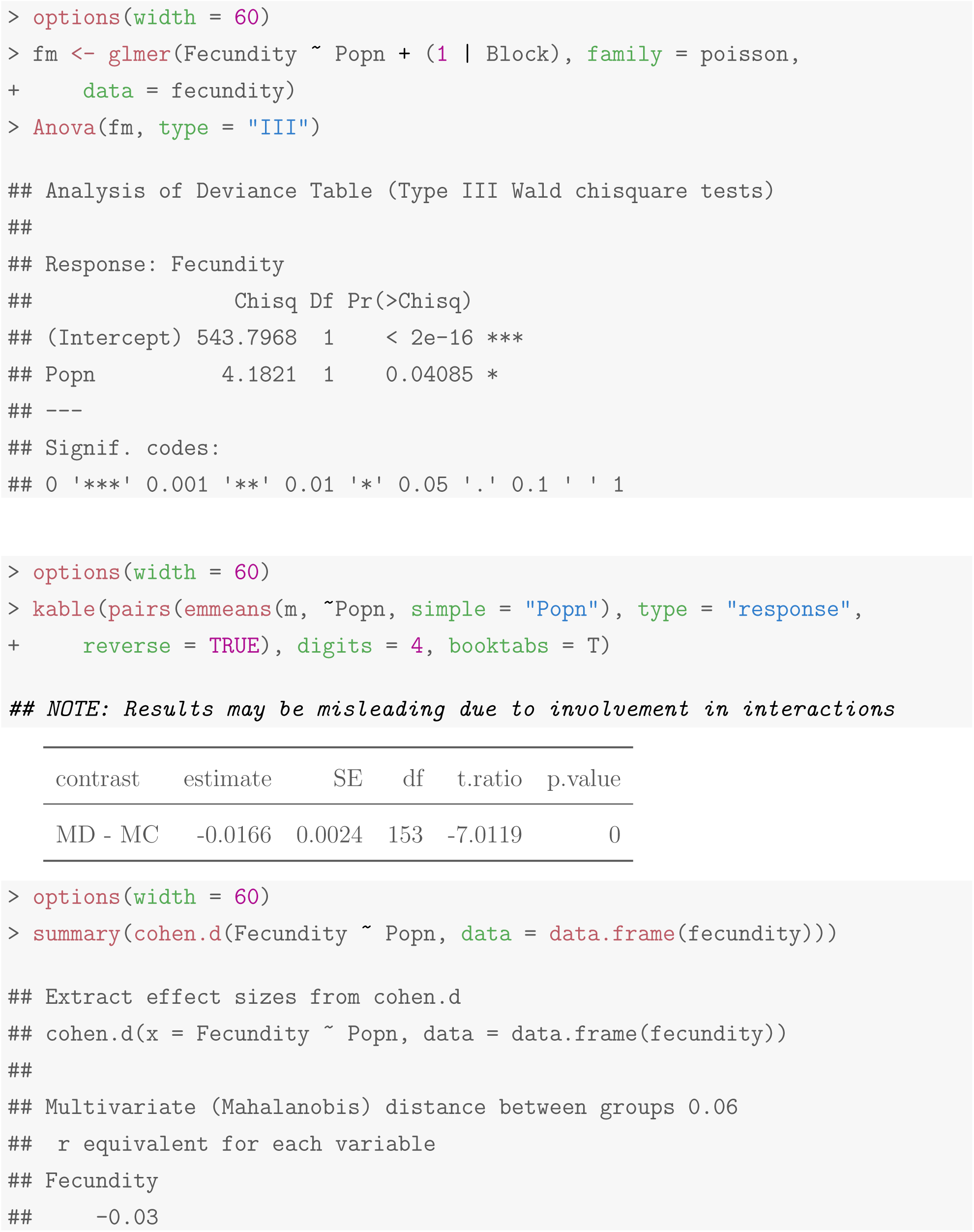

### S1.3 Time to eclosion

#### S1.3.1 All four blocks

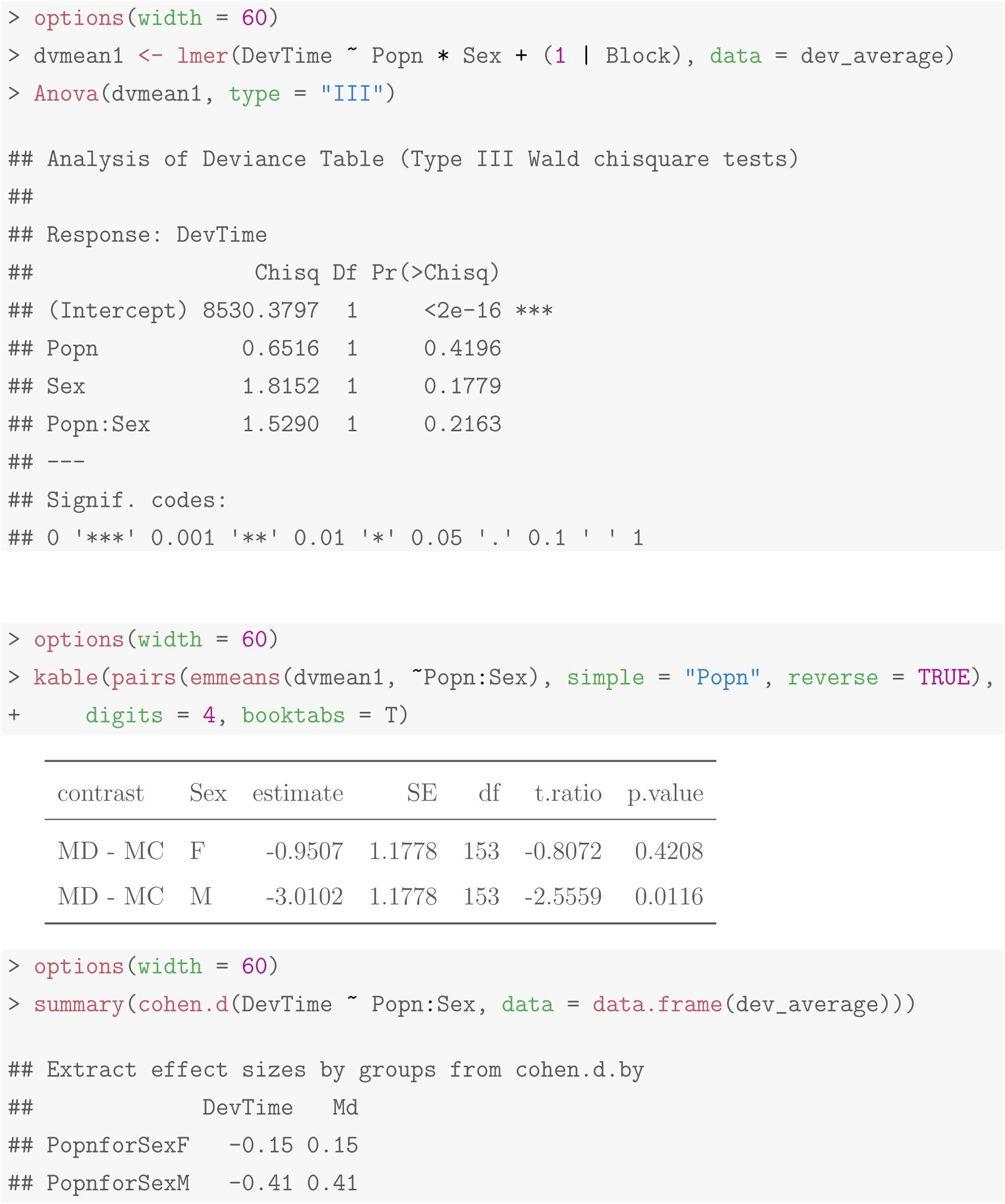

#### S1.3.2 Excluding block 4 data

Visual inspection of the data indicated that block 4 could be an outlier. The analyses were therefore repeated, now excluding data from block 4.

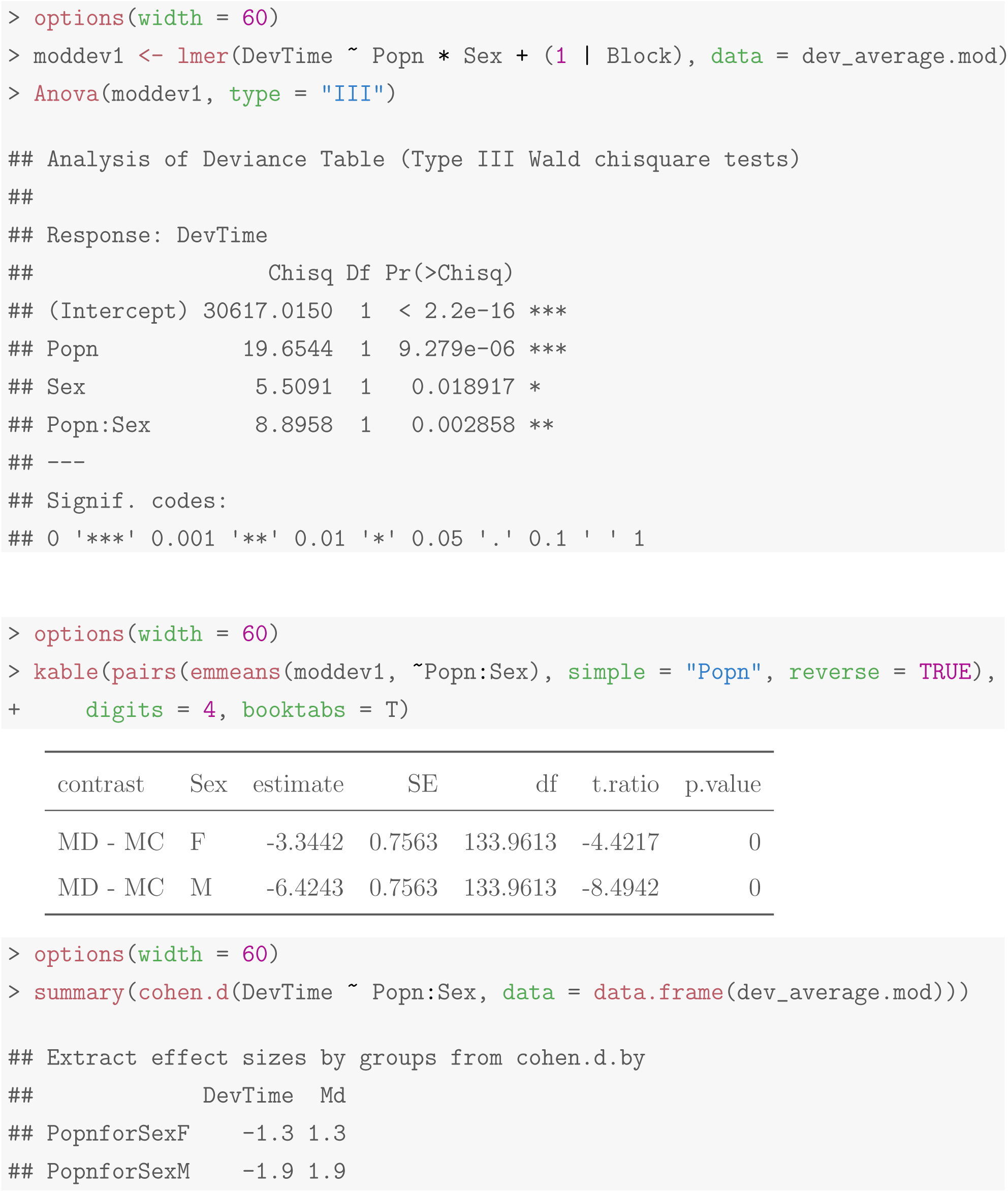

### S1.4 Time to death by desiccation

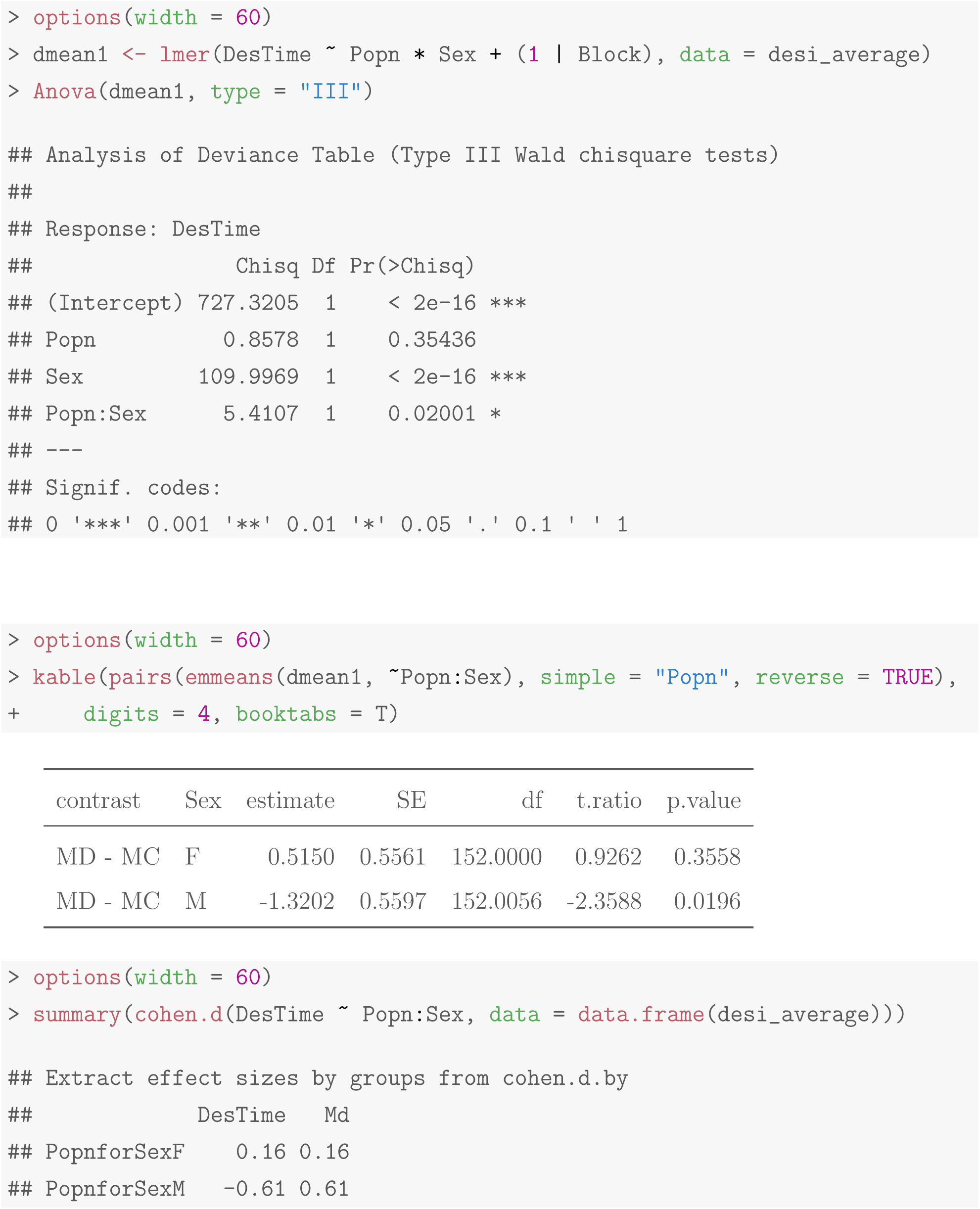

## S2 Behavioral traits

### S2.1 Locomotor activity

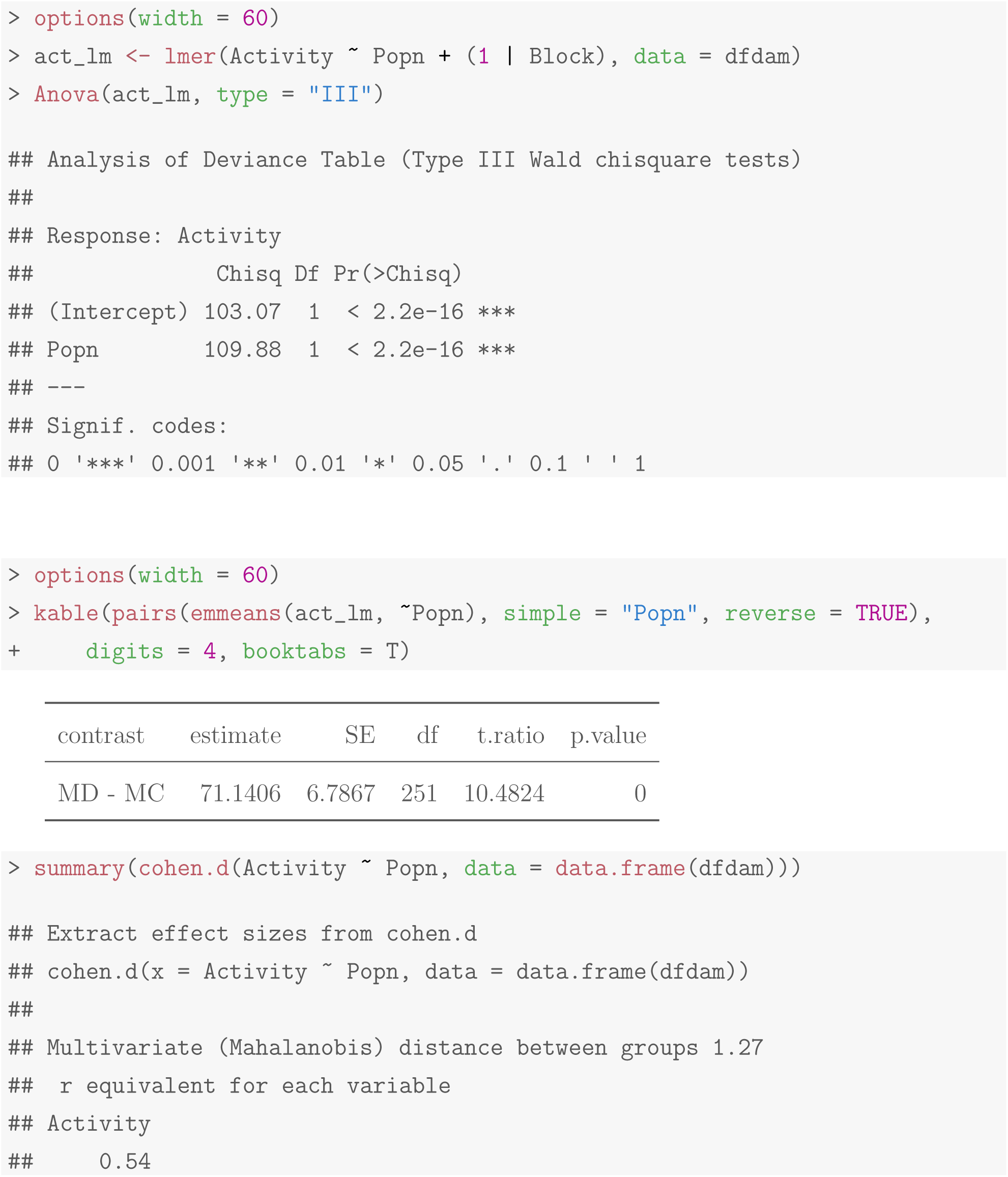

### S2.2 Rest duration

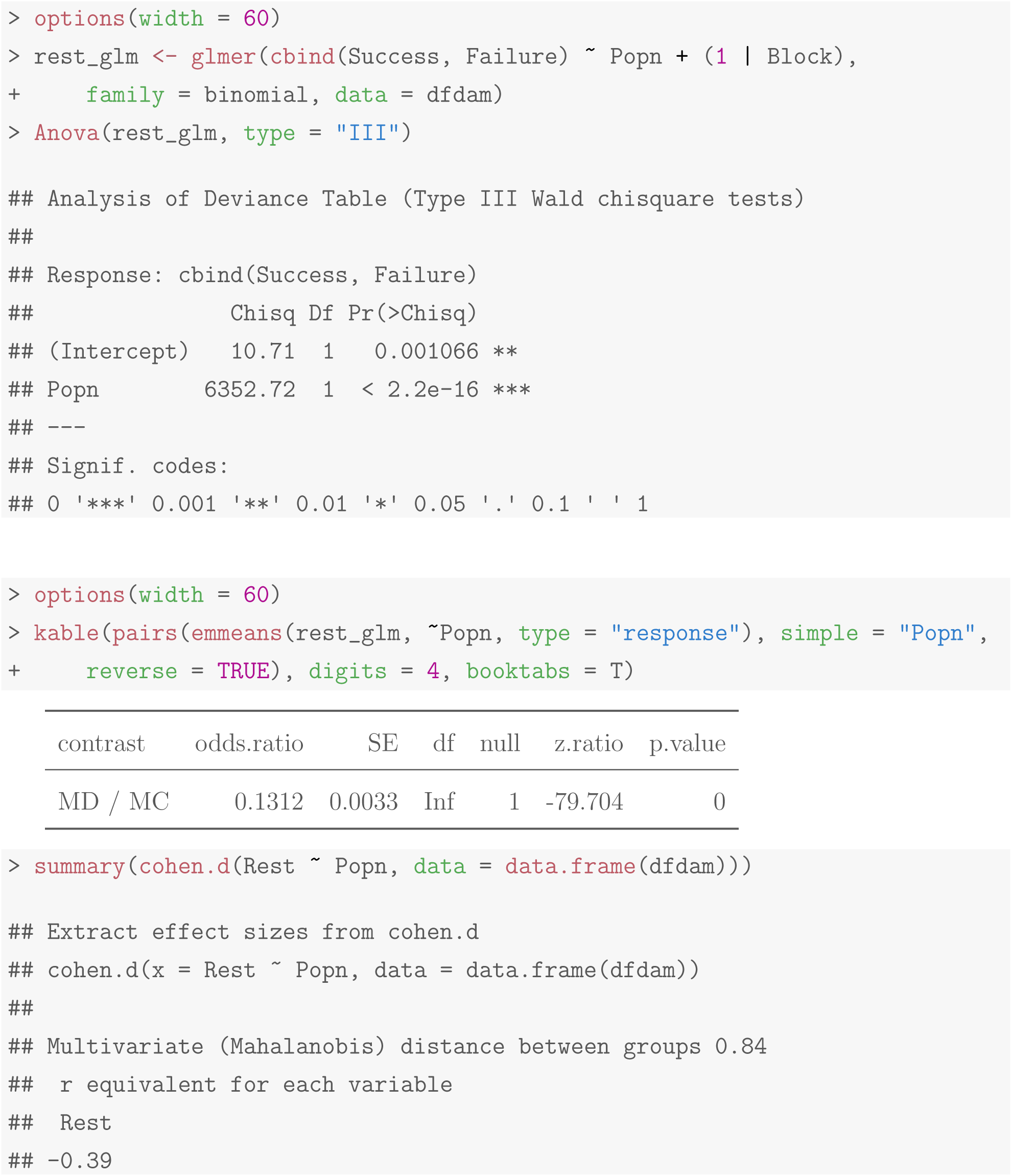

### S2.3 Mating latency

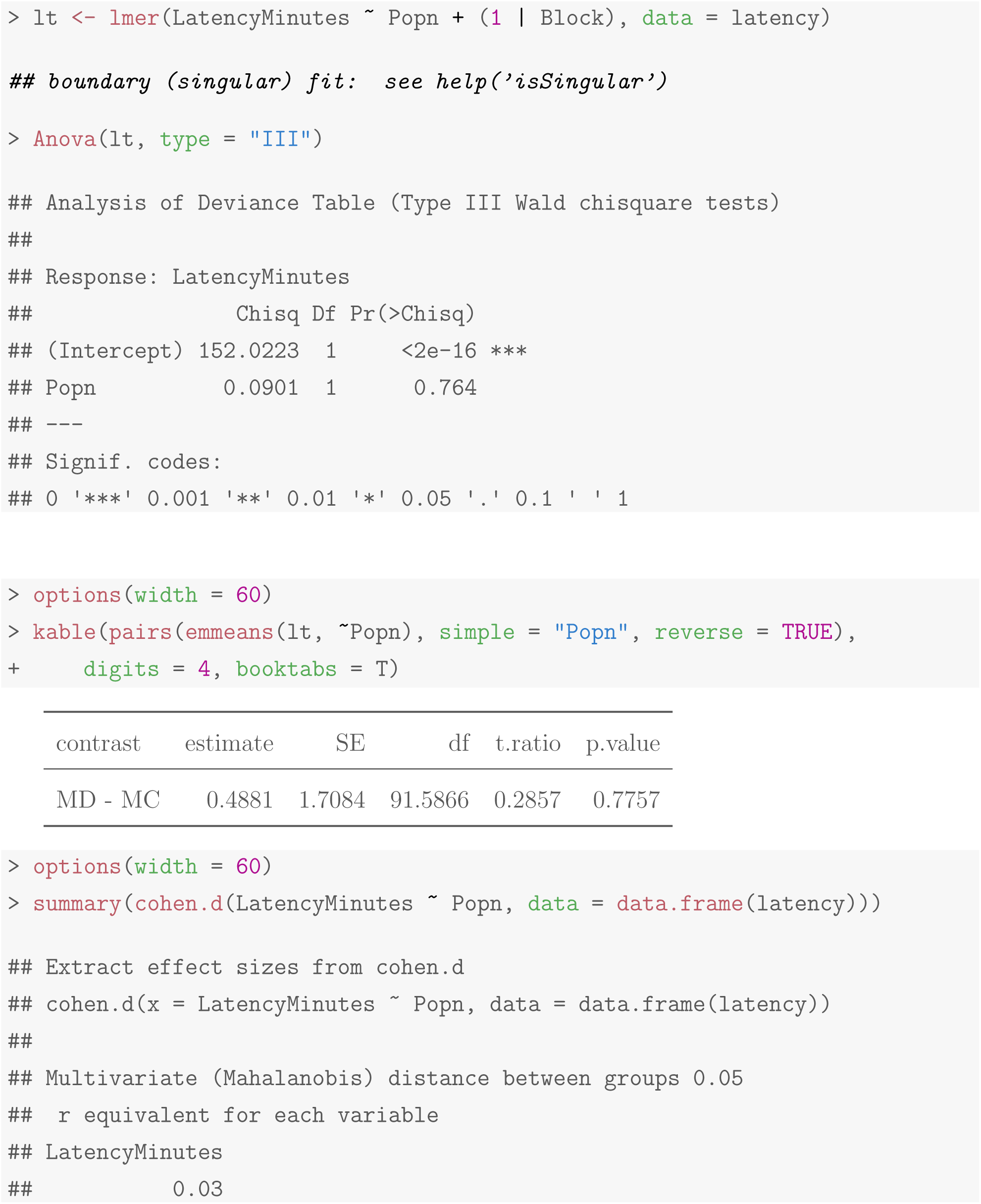

### S2.4 Copulation duration

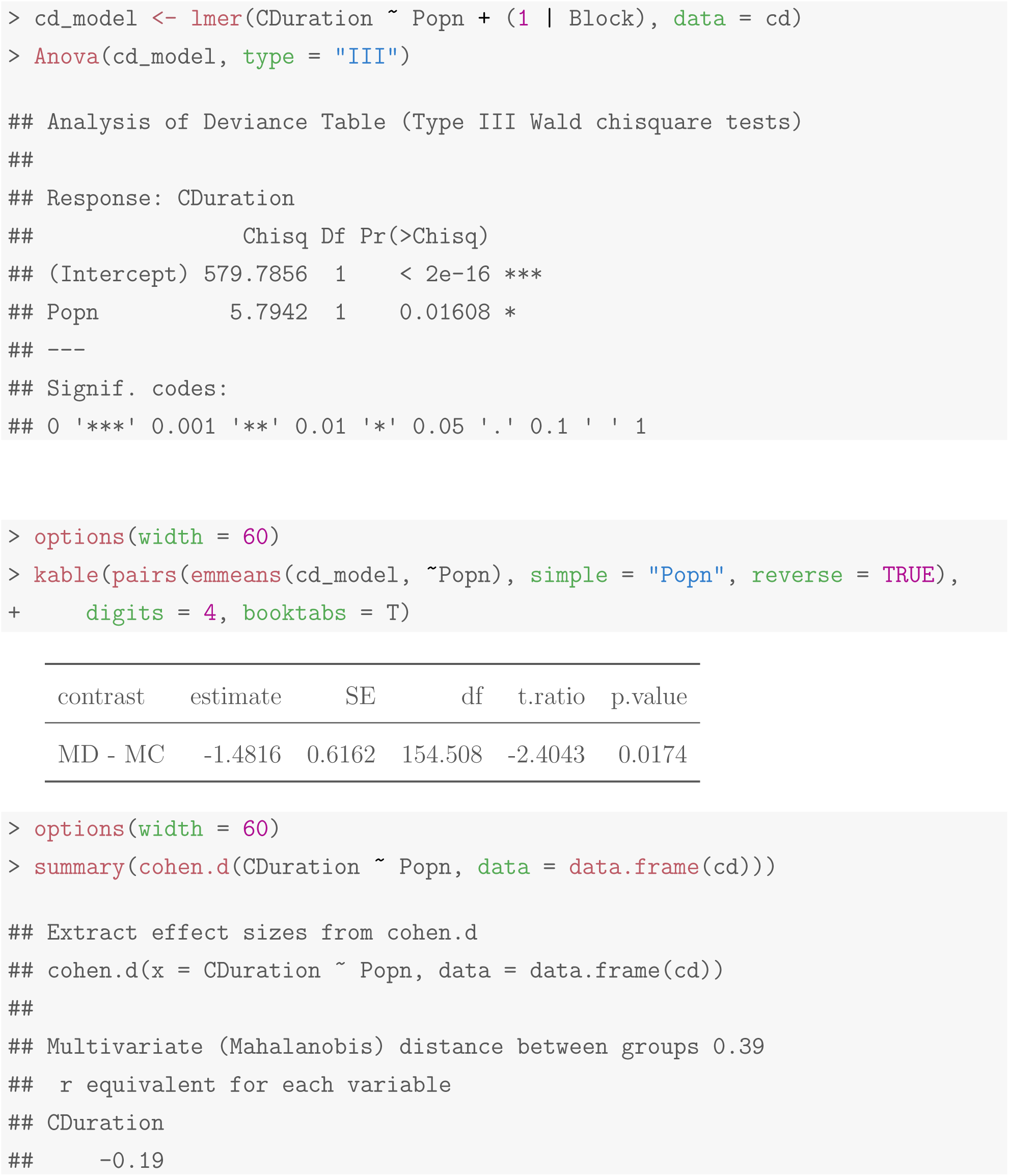

### S2.5 Mating properties

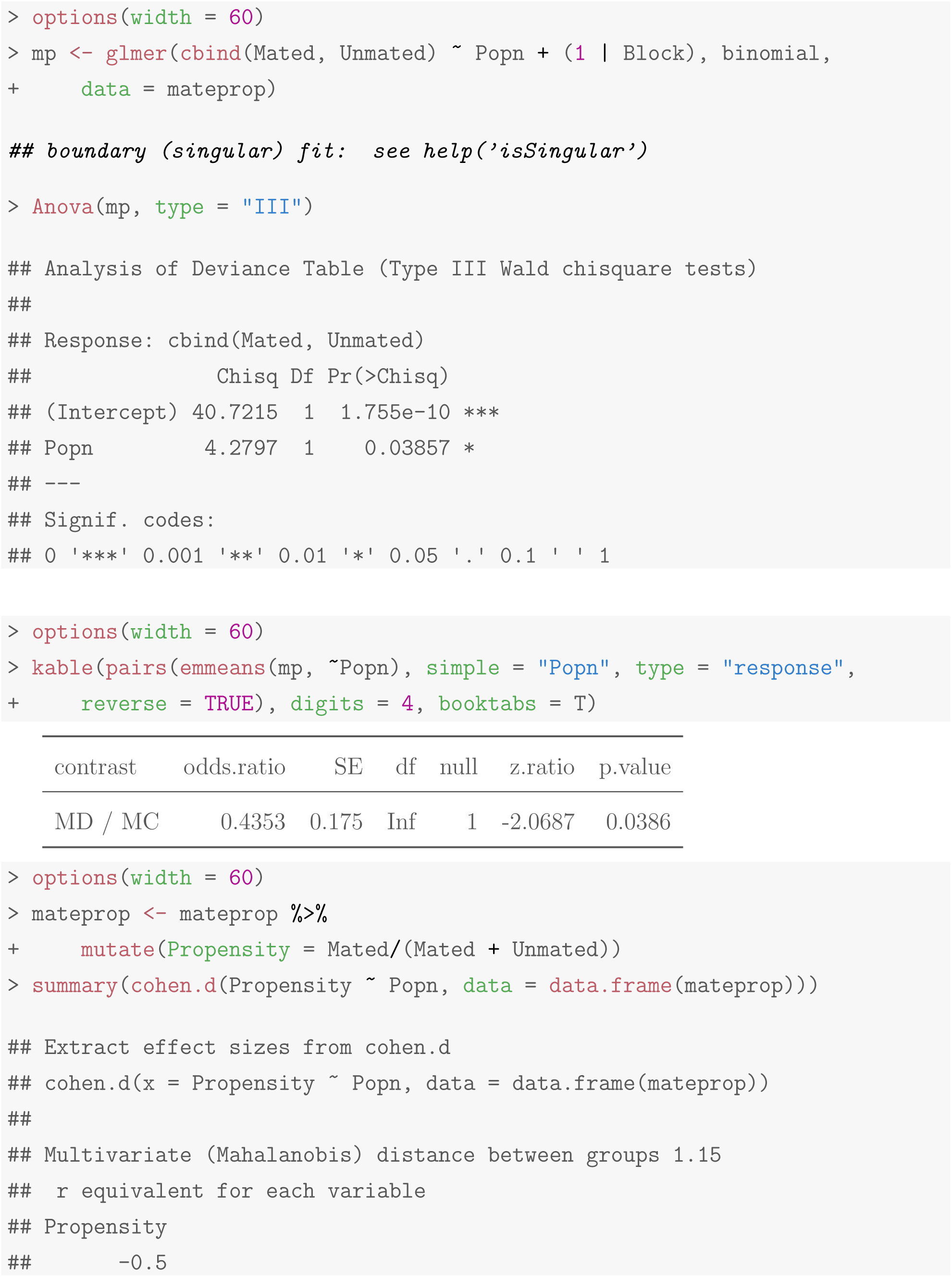

### S2.6 Male-male aggression

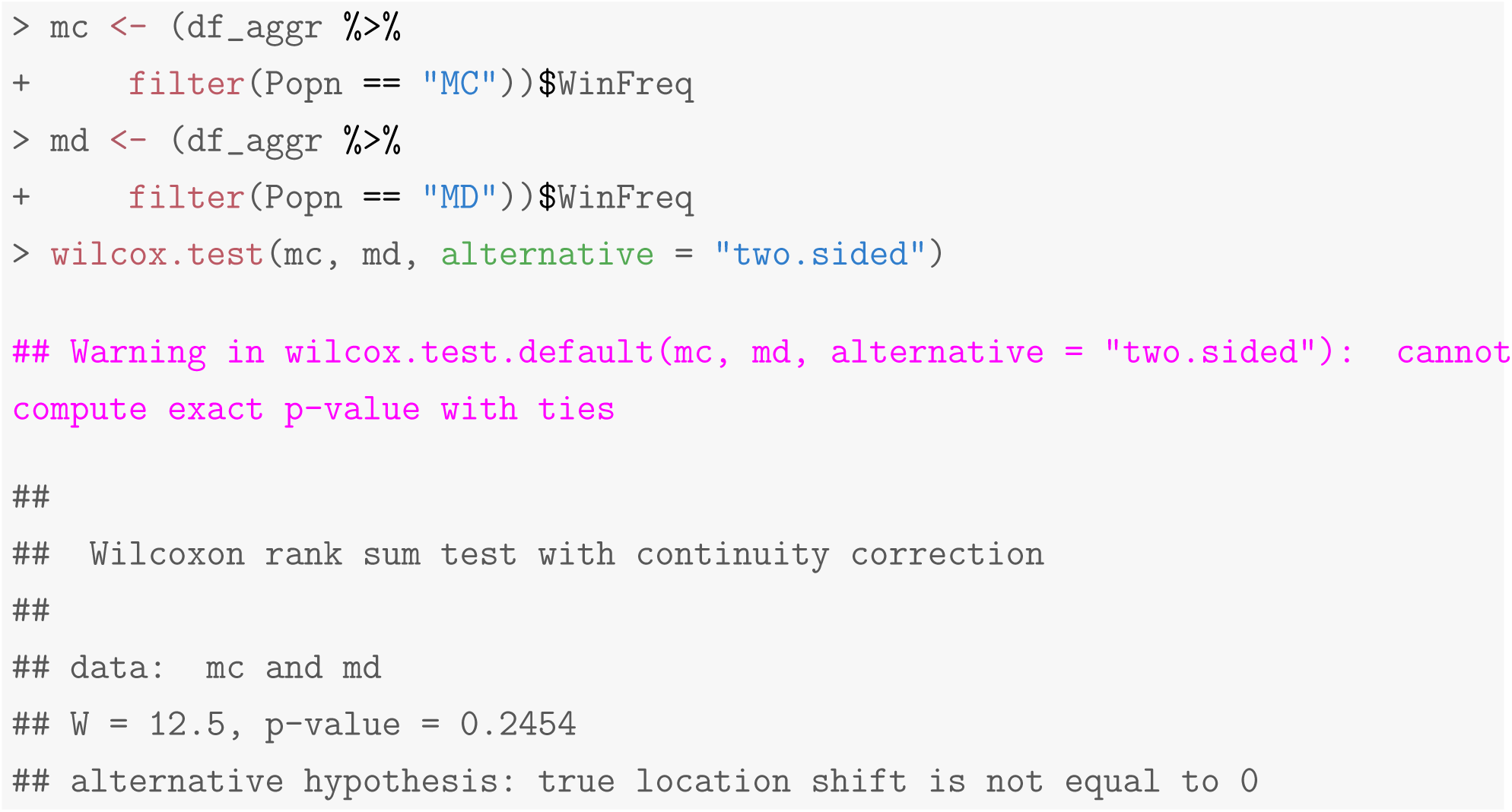

## S2 Text for “Costs of dispersal evolution under larval malnutrition”

### Recipe for modified fly food

We use a modified banana-jaggery medium with the amount of dry yeast powder reduced to one-third of that of the standard food. The banana-jaggery medium is a complex form of fly feed, of which most constituent parts contribute multiple kinds of complex sugars, lipids and other micronutrients (Nutrition, 1977). Unlike other simplified media (Lee and Micchelli, 2013), it is not possible to isolate a single source of carbohydrates or protein whose levels can be changed in a factorial design to produce multiple kinds of malnutrition regimes. Yeast powder is by far the simplest and most standardized component of the food as it is obtained from the same source without the kind of batch-to-batch variation that is inevitable in natural components like bananas or jaggery. Since yeast is the primary source of protein in the banana-jaggery recipe, its reduction corresponds to reduction in the amount of total available protein by about 20%, with some auxiliary reduction in carbohydrates. Standardisation experiments showed a clear increase in time to eclosion as well as reduction in average dry body weight in both sexes (unpublished data) in unselected flies subjected to this malnutrition regime of one-third yeast powder, suggesting that it is strong enough to impose some amount of selective pressure over several generations.

**Supplementary Table S21:**
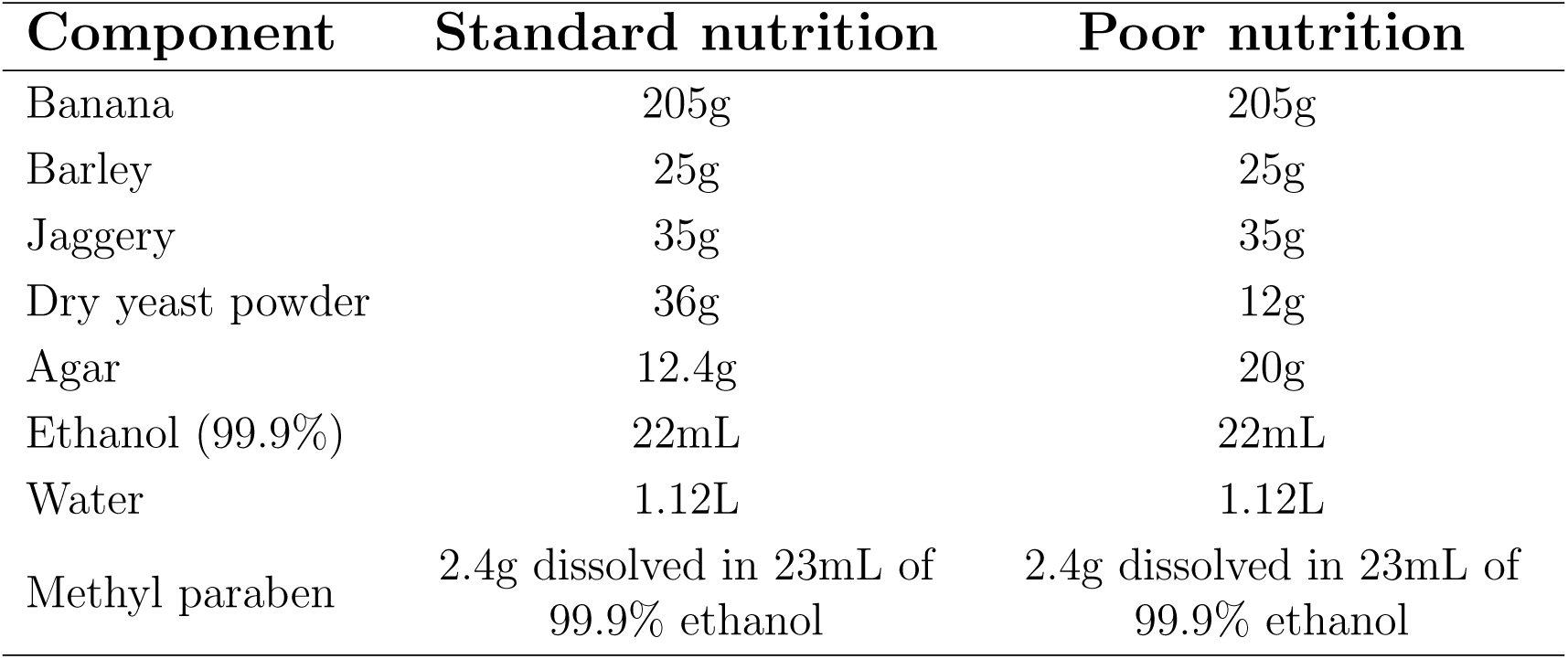
Recipes of banana-jaggery medium used for stock rearing and experiments. The detailed procedure for food preparation is available at request.

## Notes

Funding: This project was supported by grant # CRG/2018/001333 from the Science and Engineering Research Board, Department of Science and Technology (DST), Government of India, and internal funding from the Indian Institute of Science Education and Research (IISER), Pune. BV was supported by IISER Pune. AK was supported by IISER Pune and the INSPIRE Fellowship from the DST. AM was supported by a senior research fellowship from the Council of Scientific and Industrial Research (CSIR), Government of India.

### Competing Interest Statement

The authors have declared no competing interest.

